# Predictive coding in the auditory brainstem

**DOI:** 10.1101/2023.12.31.573202

**Authors:** Alain de Cheveigné

**Affiliations:** Laboratoire des Systèmes Perceptifs, Centre National de la Recherche Scientifique, École normale supérieure (Paris), and University College London

## Abstract

Predictive coding is an influential concept in sensory and cognitive neuroscience. It is often understood as involving top-down prediction of bottom-up sensory patterns, but the term also applies to feed-forward predictive mechanisms, for example in the retina. Here, I discuss a recent model of low-level predictive processing in the auditory brainstem that is of the feed-forward flavor. Past sensory input from the cochlea is delayed and compared with the current input, with a delay that is tuned with the objective of minimizing prediction error. This operation can be performed uniformly across peripheral channels, or independently within each peripheral channel with parameters local to that channel. The result is a sensory representation that is invariant to a certain class of masking sounds (harmonic, quasi-harmonic, or spectrally sparse), thus contributing to Auditory Scene Analysis. The purpose of this paper is to discuss that model in the light of predictive coding, and examine how it might fit within a wider hierarchical model that supports the perceptual representation of objects and events in the world.

## I. Introduction

In predictive coding, an incoming sensory pattern is compared with a predicted pattern, and the outcome of this comparison – the error – serves both to tune the predictive model, and as an input for the next stage of processing. Predictive coding is an influential idea in sensory and cognitive neuroscience (Attneave 1954, Barlow 1961, Srinivasan et al 1982, Rao and Ballard 1999, Clark 2013; Friston 2018). A perfectly predictable pattern carries no information beyond that carried by the predictor (it is redundant). Even if the prediction is imperfect, the error may be less costly to represent and/or easier to process and/or more informative than the raw input. Furthermore, the *parameters* of the predictive model, fit to the stimulus, characterize the stimulus, while the model itself, tuned over a longer time scale, captures the statistics of the sensory environment. Predictive coding is one of a constellation of related ideas, including unconscious inference (Alhacen 1030; Helmholtz 1867), efficient coding (Shannon 1948), epistemological automaton (MacKay 1956), redundancy reduction (Barlow 1961), forward models in motor control (Wolpert et al 1995), reverse hierarchy (Hochstein and Ahissar 2002), efferent copy (Crapse 2008), compression (Schmidhuber, 2009), homeostasis (Friston 2010), Bayesian inference (Aitchison and Lengyel 2017), and so on. In each case, an incoming observation is compared against an internally-generated template, and the mismatch used to adjust the model and remove redundancies.

However, “predictive coding” covers two quite different concepts (Spratling 2015). Given a hierarchy of processing levels, prediction can occur in the *ascending* direction (bottom-up Fig. 1 A) or *descending* direction (Fig. 1 B). An example of the former is center-surround receptive fields in the retina (Srinivasan et al 1982; Hosoya et al 2005), an example of the latter is the recurrent processing proposed for the visual cortex by Rao and Ballard (1999). Top-down prediction is usually assumed to operate within a *hierarchy* of predictive stages ordered from concrete to abstract (Fig. 1C) (e.g., Rao and Ballard 1999; Spratling 2017; Friston 2018), with messages exchanged in both directions. Bottom-up prediction is embodied in the Linear Predictive Coding (LPC) method popular for spectral estimation in speech processing (Markel and Gray 1976), while top-down prediction is related to Bayesian models of perception that combine a prior (parameters of the predictive model) and evidence (the sensory pattern) to infer the causes of sensory input. Both flavors of predictive coding could play a role within the auditory system (Heilbron and Chait 2018).

**Fig. 1.**
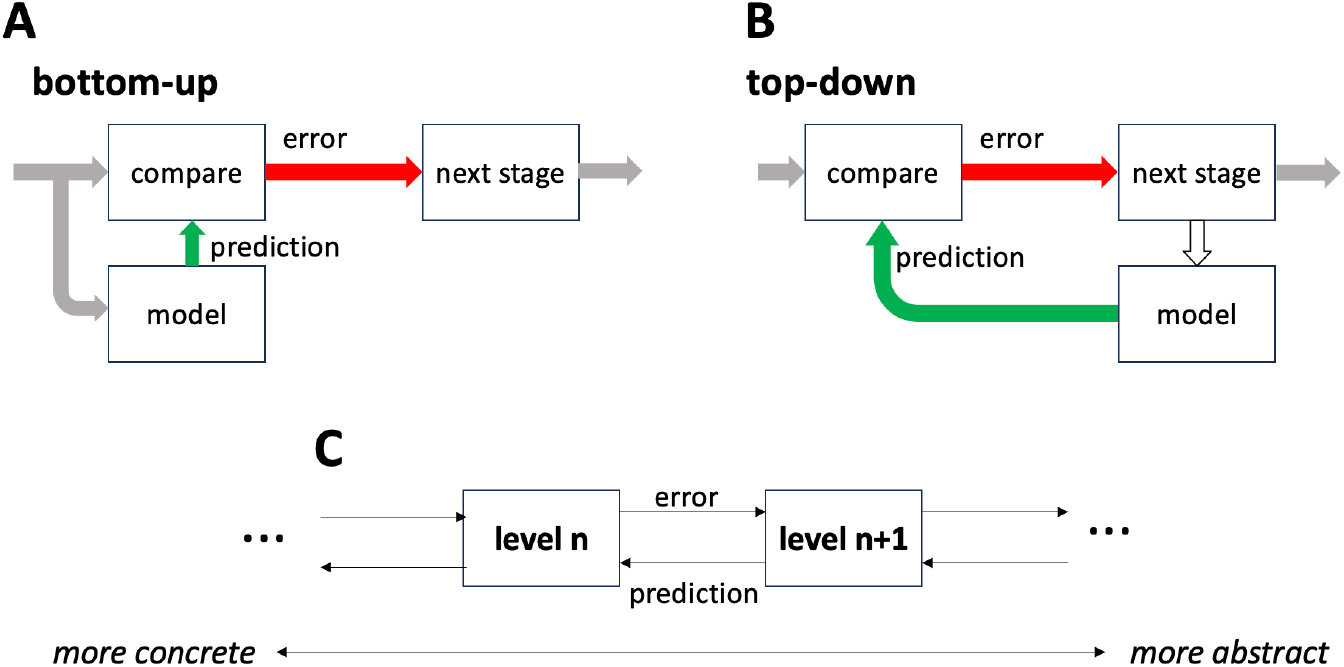
Two forms of predictive coding. A: The prediction is produced by a feed-forward model (bottom-up) fed from incoming patterns. B: The prediction is produced by a feed-back model (top-down) fed from a higher-level representation. C: Hierarchy of predictive coding stages ordered from concrete to abstract.

Here, I reinterpret a recent model of low-level auditory processing as a form of predictive coding of the bottom-up flavor: low-level sensory input is compared, at each instant, with a prediction derived from its previous values. I then take advantage of this concrete, bottom-up model to explore issues that arise when considering the alternative hypothesis of top-down predictive coding, and discuss how the two approaches might complement each other.

## II. The in-channel cancellation model

*In-channel cancellation* (ICC) is a hypothetical processing mechanism for auditory scene analysis, to help address a listener’s need to handle “cocktail party” situations involving complex acoustic scenes with multiple concurrent speakers or polyphonic music. In particular, it accounts for our ability to ignore masking sounds that are *harmonic, quasi-harmonic*, or *spectrally sparse*. The structure and principle of the ICC model is outlined here, more details and references are provided elsewhere (de Cheveigné 2023a).

Patterns of acoustic vibration are transduced within the cochlea and carried by an array of auditory nerve fibers to the auditory brainstem (Palmer and Rees 2010), an auditory analog of the retina in terms of level of processing (Sitko and Goodrich 2021). The cochlea implements a bank of frequency-selective filters, with each channel transduced by one of ~3000 hair cells into a pattern of spikes carried by a set of ~10 fibers (in human). In the ICC model, spikes within a channel feed a “neural cancellation filter” that consists of a gating neuron with one excitatory synapse fed directly, and one inhibitory synapse fed via an adjustable delay of duration *τ*_*k*_ where *k* is the index of the channel (Fig. 2A, red). The properties of this neuron are assumed to be such that every direct spike is transmitted *unless* a spike arrives simultaneously on the delayed pathway, within some short time window of coincidence. The effect is to modify the statistics of the spike train, depleting it of inter-spike intervals of duration close to *τ*_*k*_.

**Fig. 2.**
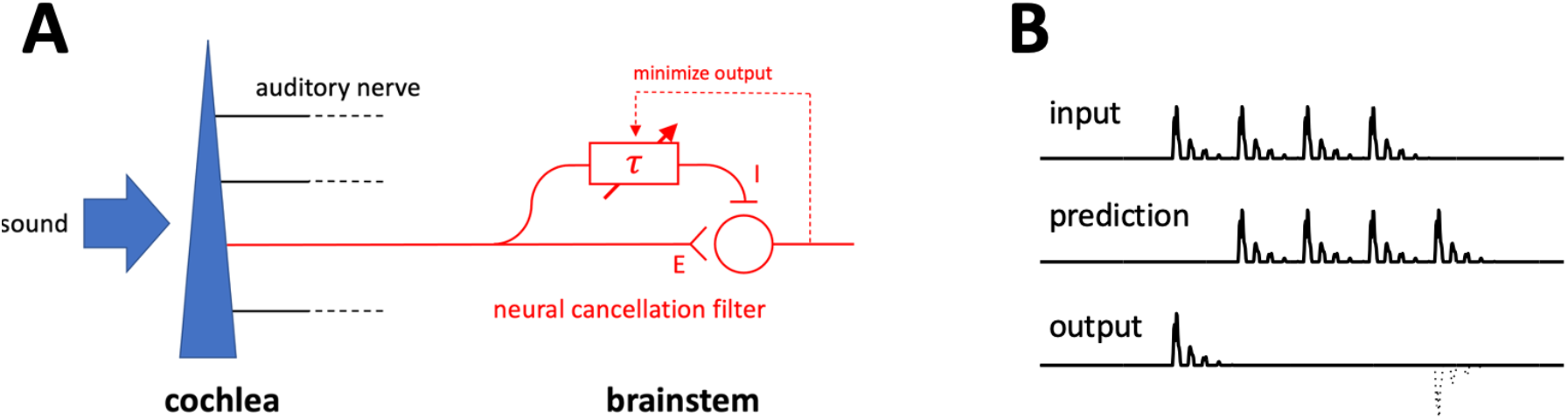
A: Schematic representation of the cochlea as a bank of bandpass filters followed by a neural cancellation filter (one channel shown). The neural cancellation filter depletes the auditory-nerve spike train of intervals equal to *τ*, thus attenuating correlates of sound components with that period. B: instantaneous spike probability at the excitatory synapse (top), inhibitory synapse (middle) and at the output of the gating neuron (bottom) in response to a short periodic stimulus. Only the first period reaches the output.

The delay parameter *τ*_*k*_ is determined automatically based on a criterion of minimum output, and it tends to lock on to the period (or pseudo period) of the signal that dominates the channel. In the presence of a masking sound that is *harmonic, quasi-harmonic*, or *spectrally sparse*, the filter cancels the correlates of that sound, potentially allowing components of a weaker concurrent sound to emerge. An earlier model (harmonic cancellation, HC, de Cheveigné 1993; 2021) used the same delay *τ* in all channels, making it effective only for the narrower class of harmonic maskers, whereas the ICC model is more flexible. Together, the HC and ICC models can account for a range of psychophysical results that involve masking with such sounds (as reviewed by de Cheveigné 2021, 2023a). The neural circuitry required by these models (fast excitatory-inhibitory interaction) is similar to that hypothesized by the classic equalization-cancellation (EC) model of binaural unmasking (Durlach 1963), and the within-channel version of that model (Culling and Summerfield 1995), for which physiological support has been found within the lateral superior olive (LSO) (Biederbeck 2018; Franken et al 2021).

In a linear approximation, the effects of this neural filter can be described by a delay-and-subtract filter of impulse response *h*(*t*) = *δ*(*t* − *τ*_*k*_) − *δ*(*t*), where *δ*(*t*) is the Dirac function, followed by a half-wave rectifier that embodies the fact that probability is non-negative (de Cheveigné 1993, 2023a). If the delay equals the period of a sound, the response to that sound is suppressed, as illustrated in Fig. 2B: the predicted signal (middle) is subtracted from the input signal (top), suppressing all but the first period (bottom). In the frequency domain, the effect is to suppress all harmonics of 1/*τ* (Fig. 3A, top). The cascade of cochlear and cancellation filters has a frequency characteristic that attenuates both power that is distant from the characteristic frequency (CF) *and* power that is harmonic with period *τ* (Fig. 3A, bottom).

**Fig. 3.**
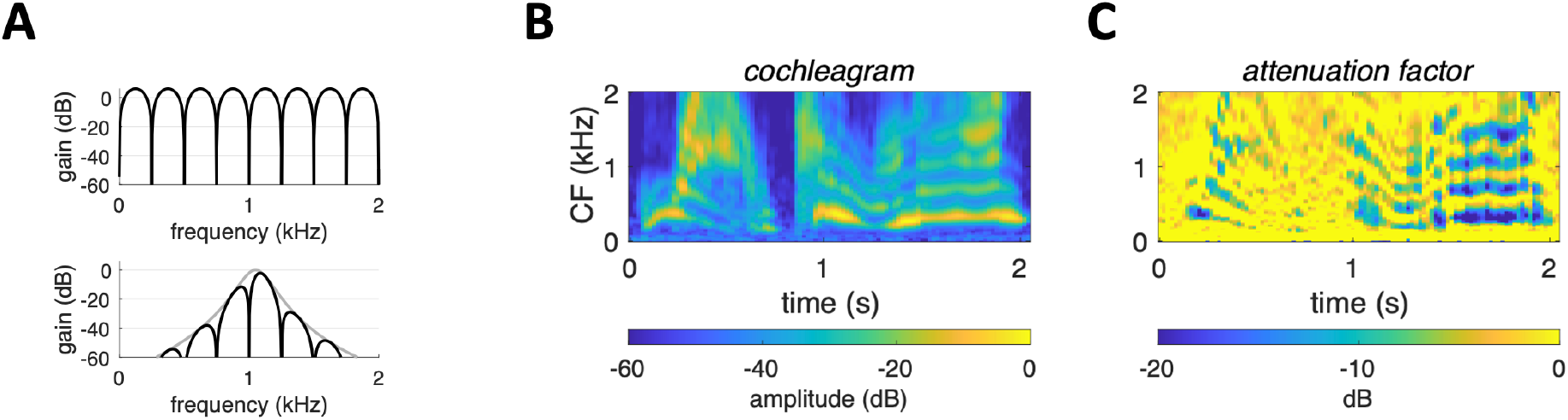
A top: transfer function of delay-and-subtract filter for *τ*=2.5 ms, bottom: transfer function of cascade of this filter with a cochlear filter with CF=1.05 kHz. B: Cochlear excitation pattern in response to a short snippet of speech (“wow, cool!”). C: Attenuation factor of the cancellation filter array at each time and CF for the speech snippet in A.

As an example, Fig. 3B shows a cochleagram, a time-frequency representation based on a modeled cochlear filterbank (Slaney 1983) in response to a short snippet of speech (“wow cool”, male voice). Panel C shows the gain of the time-varying cancellation filters applied to the output of the filterbank, also as a function of channel and time. The gain is reduced (by as much as −20dB) within the time-frequency pixels corresponding to high-amplitude components of this sound. Thus, if this speech sound were to mask a weaker target, that target might be detected or partially recovered by applying this array of time-varying filters to suppress the masker. This suggests that this hypothetical mechanism might effectively deal with highly non-stationary “real-world” sounds such as speech.

## III. ICC as predictive coding?

### A. Bottom-up

The structure of the ICC model (Fig. 2) evokes predictive coding of the bottom-up flavor, similar to the model of Srinivasan et al (1982) in which the center of a retinal patch is predicted from its neighbors and/or previous values. Here, the incoming auditory nerve spike probability is predicted from its past values (Fig. 2, left). The model is also reminiscent of LPC (Gray and Markel 1996) in which each new sample of a sampled representation is predicted as a linear combination of *N* previous samples (where *N* is the order of the LPC model). The ICC model achieves both *redundancy reduction*, in that the cancellation filter strips the representation of redundant, periodic patterns (Fig. 2B), and *redundancy exploitation* (Attneave 1954; Barlow 1961), in that the optimal delay is the period of the sound, predictive of its pitch (de Cheveigné 1998; 2023). The period is an *abstract* quantity and, in this sense, ICC may contribute to the concrete-to-abstract hierarchy depicted in Fig. 1C (see also Sect. IV, B).

### B. Top-down

There is a very different class of models in which incoming patterns are predicted from abstract internal models, according to a principle akin to “analysis by synthesis” (Helmholtz 1867; Braitenberg 1984; Kawato et al 1993; Friston 2002, 2018; Spratling 2017). There is keen interest in such top-down models, in particular due to theoretical properties of certain formulations (Friston 2010, 2018; Friston et al 2009; Bastos et al 2013; see also Sims 2017; Litwin and Miłkowski 2020; Andrews 2021; Williams 2021; Hemmatian 2023). Re-casting ICC as a top-down predictive model might allow us to “hitch onto that wagon” and reap its theoretical benefits. An additional motivation is the need to explain massive efferent pathways that link most stages of the auditory system between cortex and cochlea (Schofield 2012; Romero and Trussell 2022). Together, these reasons encourage us to consider whether it is possible to translate, or insert, ICC into a top-down model. Here I review three simple schemes, numbered for easy reference.

1. Splitting the ICC model of Fig. 2A in two, parameters *τ*_*k*_ can be treated as message from a higher level that evaluates the filter output, to a lower level that implements the filter. This reformulation makes the model seem “top-down”, but with little added value.
2. The predictive filter of Fig. 4A can be modified by deriving the “prediction” from its output (Fig. 4B) rather than input, thus making it “top-down.” A moment’s thought tells us this is a bad idea: if prediction were effective, i.e., the error was zero, there would be nothing to feed the prediction. If we persisted, we would discover that the filter has an infinite, non-decreasing impulse response (Fig. 5B, bottom). This could be alleviated by introducing a gain factor in the feedback loop, with a value less than one, but the impulse response would still be long relative to that in Fig. 5A for no clear benefit.
3. A third scheme, inspired from the concept of “analysis by synthesis” is to derive the prediction via a generative model fed from a higher, abstract stage (Fig. 5C). This too cannot work, because the temporal (or phase) information needed to correctly align the prediction with the incoming pattern is not available. As pointed out by Mumford (1992), the higher, abstract stage speaks a different “language” from the lower concrete layer. Abstraction is a many-to-one transform, not invertible, and thus an abstract trace cannot translate unambiguously to a concrete one.

**Fig. 4.**
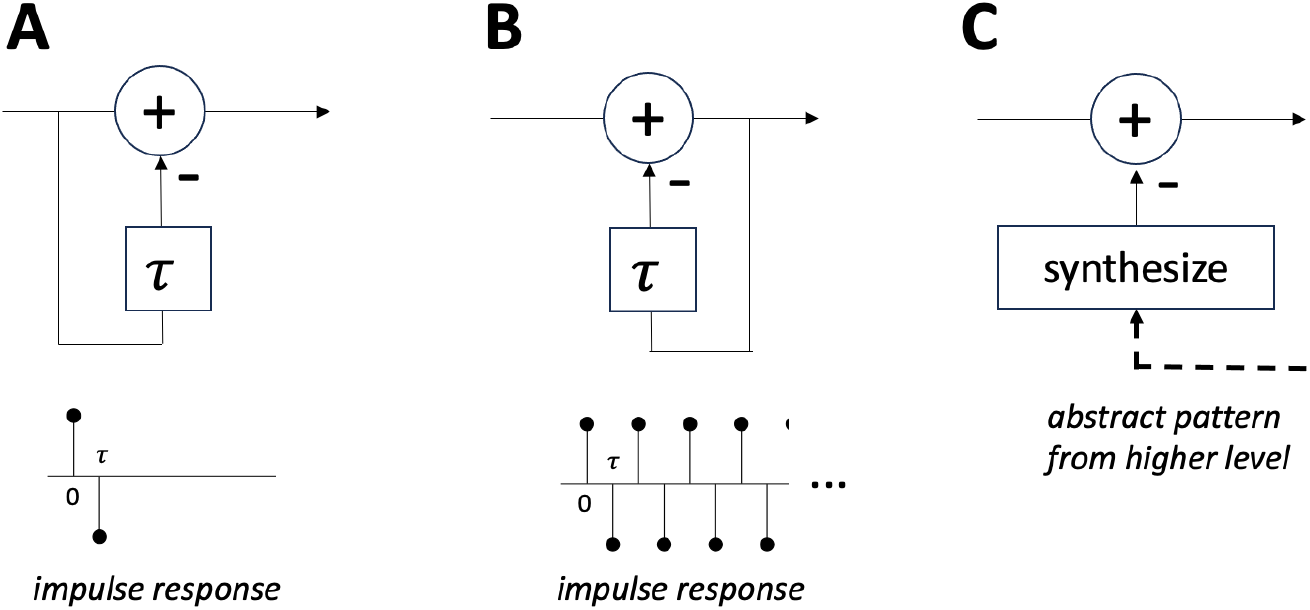
A: The bottom-up predictive coding model embodied by ICC implements a filter with a very compact impulse response that suppresses periodic input. B: Deriving the prediction from from the output leads to a filter with an infinite, non-decreasing impulse response. C: Top-down prediction from an abstract trace cannot work because temporal (or phase) information to align it with the input is lacking.

**Fig. 5.**
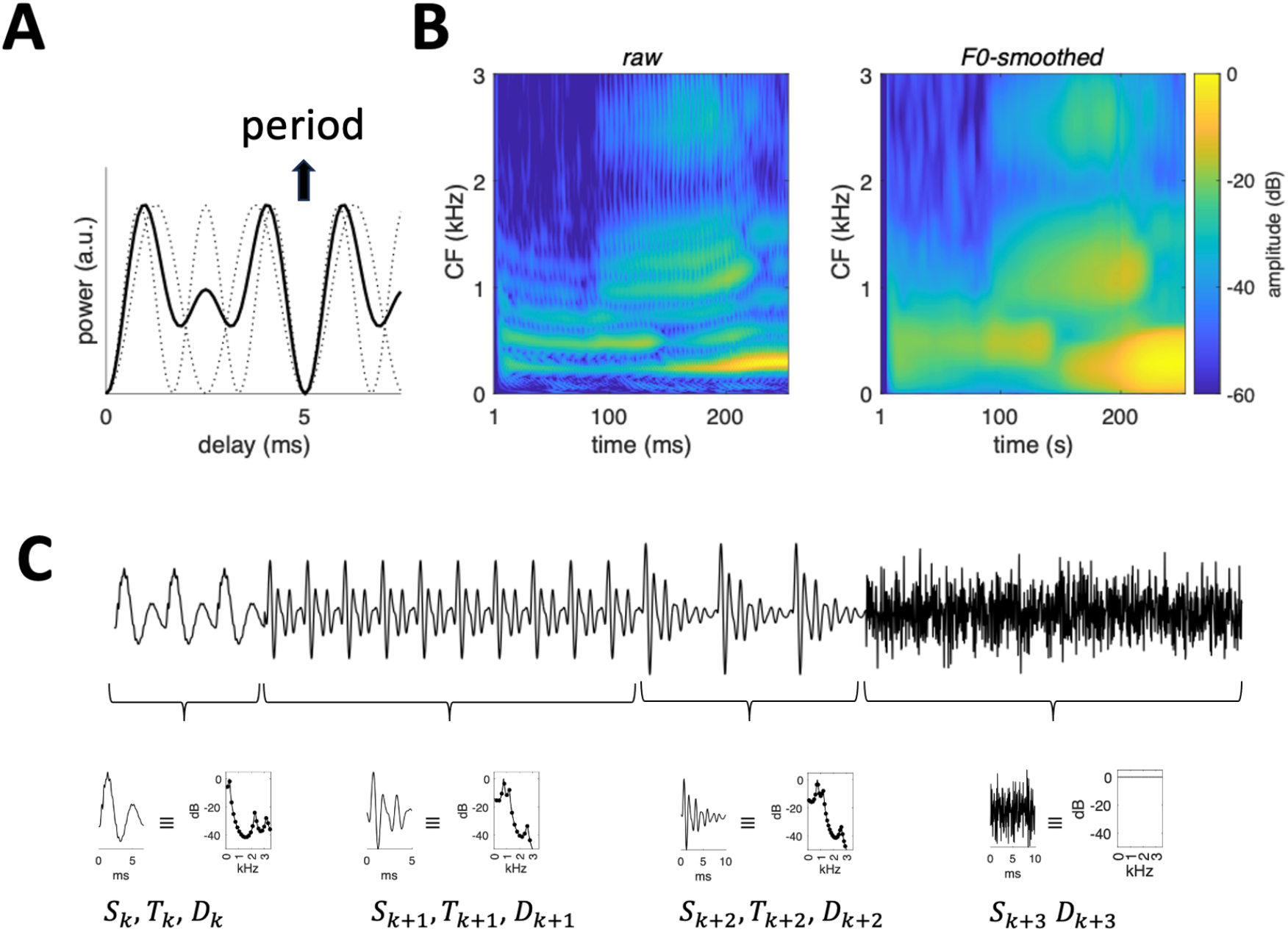
ICC as abstraction. A: The period of a 200 Hz complex tone is estimated as the first non-zero lag that yields zero output, common over channels. The black line is for CF=500 Hz (this channel responds to both harmonics 2 and 3), the dotted lines are for CF=400 Hz (harmonic 2) and CF=600 Hz (harmonic 3). B: Optimal smoothing of a time-frequency representation. Left: raw cochleagram, right: cochleagram smoothed in time (kernel duration *T* = 1/*F*_0_) and frequency (kernel width *F*_0_ = 1/*T*). C: Chunking for redundancy reduction. Each stationary segment might be coded by the *waveform during one period* (or its Fourier transform), the value of the *period* (for periodic sounds) and the *duration* of the segment.

#### Interim summary

The ICC model looks like a text-book example of bottom-up predictive coding, but straightforward attempts to reformulate it as a top-down model are not fruitful. The rest of this paper tries to better understand why, and salvage the idea in view of its multiple benefits invoked at the beginning of this section.

## IV. Let’s try again

### A. Top-down predic3ve coding revisited

Theories of perception mostly agree on a hierarchy of representations going from concrete (sensory) to abstract (perceptual, or conceptual), with messages passed between levels in the ascending and (for predictive coding) descending directions (Fig. 1C). As pointed out by Mumford (1992), messages passed between different levels of abstraction require *translation*, which could occur *before* the message is sent, or *after* it is received. Referring to Fig. 1C, the ascending error might be translated at either level *n* or level *n* + 1, and the same options are available for the descending prediction, implying four different interpretations of Fig. 1C, each plausible but with different implications and constraints. This is an example of questions that may arise when trying to make sense of predictive coding.

Top-down translation presents two difficulties. One is the difference in representation format (e.g., a retinotopic sensory pattern in a sensory area versus an abstract representation within a face area), the other is that abstraction is not invertible. For the first, Mumford (1992) suggested that a “template neuron” might code a detailed sensory pattern in the form of synaptic weights. Bottom-up translation would be performed by weights between axons from neurons in level *n* onto the dendrites of a template neuron in level *n* + 1, whereas top-down translation from that template neuron is performed by weights of on its axonal terminals in layer *n*. Incidentally, both sets of weights define the same “template,” which raises the question of how they are learned in sync. For the second difficulty, he proposed that a *family* of templates collectively represents the set of inputs that map to the abstract trace. Top-down translation then involves comparing all of these templates, sequentially or in parallel, to the input pattern, the error (or residual) being presumably the vector difference between input and closest template.

Mumford (1992) further suggested that each template might be “flexible”, for example defined by a mean and covariance (analogous to a Gaussian mixture model), or alternatively as a collection of extreme values (analogous to support vectors). This adds flexibility to the representation of the distribution of observations at level *n* that map to an abstract trace at level *n* + 1, easing bottom-up translation, but it is less clear how the residual is defined. Is it the difference from the mean of the nearest template? Is it the difference from the nearest “support vector”? He further suggested that observations might be *transformed* before prediction (for example by time-warping), increasing flexibility. As yet another alternative, Mumford (1992) also suggested that each template might be formed as linear combinations of basis-functions, on the lines of Poggio’s (1990) “Hyper Basis Function” theory. An abstract trace is then a distribution of sets of *coefficients* of such combinations, flexibility coming from the spread of this distribution rather than from the diversity of a set of templates, and/or their individual flexibilities.

In a similar direction, Rao and Ballard (1999) suggested that an abstract trace be defined based on a linear combination of basis-functions *u*_*j*_, plus a stochastic noise term:

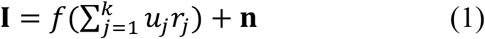

where **I** is an input that maps to the abstract trace, *f* is a function (e.g., sigmoid), *r*_*j*_ is a coefficient (“cause”) and **n** is a noise process. Here too, the abstract trace is defined as a set of coefficients, but flexibility is now determined by the noise term, rather than by the distribution of admissible sets of coefficients. Friston’s (2009) “free energy principle” requires further that this noise term be *Gaussian*: in the “Laplace approximation,” an abstract trace translates to (or is translated from) an ensemble of observations that is distributed normally in some space. The prediction residual (“error” in Fig. 1) is defined as the vector difference between observation and mean, possibly weighted by “precision” derived from the covariance matrix of the abstract trace (e.g., Friston 2002), or of the observation (e.g., Heldman and Friston 2010), or perhaps both. Redundancy reduction is achieved if fewer bits are required to represent the narrower range of the error than of the observation.

The Gaussian assumption is restrictive compared to the arbitrarily-shaped distribution of Mumford (1992), but Friston (2009, p298) argues that it can easily be satisfied by applying the appropriate *non-linear transform* to the observations. This shifts the “heavy-weight lifting” of modeling a complex distribution (e.g., visual patterns that match the concept “dog”) to that bottom-up transform. ICC might be interpreted as implementing one such transform, as it projects a diverse set of (quasi-)harmonic sounds to a space where they are distributed near zero.

Spratling (2017) pointed out the similarity between bottom-up predictive coding (e.g., LPC) and top-down predictive coding (e.g., Rao and Ballard 1999) which can both be expressed by an equation analogous to Eq. 1, in which the basis-functions *u*_*j*_ are intervals of recent observations for LPC, while for top-down predictive coding they belong to an internal generative model, fixed or slowly-varying. This opens the possibility of *hybrid* ascending and descending prediction, again suggesting a bottom-up component in top-down predictive coding.

If one insists on a “free-form” distribution (rather than Gaussian), the “error” cannot take the form of a simple difference. For a single observation, it might be quantified by a measure of *surprise* (a scalar), and for a set of observations (or an uncertain observation), by a measure such as *KL divergence, D*_*KL*_(𝒪 ∥ 𝒜), where 𝒪 represents the distribution of the observation(s) and 𝒜 that of the abstract trace (e.g., Pinchetti et al 2002). If divergence is small, the abstract trace is a good model of the observations, and so the error message can be empty. If divergence is large, a useful message might consist of the surprising observations themselves, or some transform thereof (e.g., statistic). Redundancy reduction is achieved if the latter event is rare. Note, however, that KL divergence requires *parametrization* of the distributions, which removes some (but not all) “freedom” from the free-form assumption.

It is worth noting also that KL divergence is not symmetric. *D*_*KL*_(𝒪 ∥ 𝒜) ≈ 0 means that *the observations fit the abstraction*, whereas *D*_*KL*_ (𝒜 ∥ 𝒪) ≈ 0 means that *the abstraction fits the observations*. The former might be true and the latter not if recent observations are more tightly distributed than expected, in which case a new abstraction might be considered, possibly nested within the old (e.g., a category “poodle” within the category “dog”). For this to occur, the distribution 𝒪 must be reliable, which may require gathering multiple observations over some interval of time. This, in turn, requires *chunking* the observation stream, yet another bottom-up process. This suggests a different narrative: predictive coding achieves redundancy reduction for the purpose of *efficient memory* of the stream of observations (de Cheveigné 2025).

Finally, in the “forward-reverse optics” model of Kawato et al (1993), a complex visual scene is analyzed by a combination of two processes, one top-down, the other bottom-up. The former attempts to generate retinal patterns from an internal representation of the scene, as in other analysis-by-synthesis schemes. The authors refer to it as a “direct optics” model because it mimics, within the brain, the optical pathway from scene to retina, along the lines of a forward model in motor control (Wolpert and Kawato 1998). The bottom-up process, referred to as “reverse optics,” attempts to translate retinal patterns into a representation of the scene, as in a classic computer vision system. This second, bottom-up, process does not need to perform perfectly, and indeed, it cannot, given the ambiguity of scenes with occlusion, etc. Its role is to provide an initial “guess” of the scene that allows the search space of the top-down process to be pruned. Transposing to audition, we could hypothesize that ICC contributes to a “reverse acoustics” model, applicable to parse scenes that include harmonic or quasi-harmonic sources.

In sum:

1. The term “predictive coding” has been applied to a variety of very different concepts.
2. “Analysis-by-synthesis” is problematic because abstraction is not invertible. “Prediction error” may make sense in terms of surprise or KL divergence, but not in terms of a vector *difference* unless restrictive assumptions apply. While a *property* of an observation may be predicted from an abstract trace, the value of the observation may not.
3. Top-down predictive coding relies on bottom-up processing for much of the “heavy lifting,” in particular to perform abstraction between layers of a hierarchical representation.

### B. In-channel cancellation contributes to abstraction

ICC was earlier presented as a kind of adaptive filter, the output of which has the same level of abstraction as its input (i.e., a rapidly-varying pattern of firing probability). Here, I look at how it might produce slowly-evolving abstract quantities, thus contributing to a first stage in a hierarchy of abstractions.

A first such quantity is the *period*, estimated by fitting the ICC or HC model with a criterion of minimum output (Fig. 5A). It is an abstraction in that different periodic sounds may share the same period. A wider class of sounds are *approximately periodic* in various ways: limited duration, slow amplitude or frequency changes, periodicity within a restricted spectral region, etc. (as in the speech example of Fig. 3B). For these, a measure of *periodicity* may be informative, for example the cancellation filter output/input power ratio (as plotted in Fig. 3C for individual channels over time). Period and periodicity are perceptually relevant, the former as a cue to *pitch*, the second as a possible contributor to an aspect of *timbre*.

Another abstraction is the *spectrum*, which comes in many variants according to how it is calculated. A time-frequency representation can be calculated based on the short-term Fourier transform (*spectrogram*), or as instantaneous power at the output of a cochlear filterbank (*cochleagram*, e.g. Fig. 3B), or instantaneous probability at the output of a more realistic peripheral model. Again, it is abstract in that many different sounds share the same spectrum or spectrogram, and perceptually relevant as a predictor of *timbre*, for example that of a vowel. Unfortunately, this representation is not invariant relative to the fundamental frequency *F*_0_ = 1/*T* of a sound: individual harmonics may emerge at low frequencies, and a pulsed temporal structure may emerge at high frequencies (Fig. 5B, left), whereas timbre is usually thought to reflect the smooth time-frequency *envelope*. Here again, the ICC/HC model contributes by allowing *period-adaptive smoothing* to approximate that envelope (Fig. 5B, right). This is the principle of the well-known STRAIGHT method of spectro-temporal estimation (Kawahara et al 1999). It offers yet another level of abstraction, as sounds with different spectra/spectrograms (due to different fundamental frequency) map to the same smooth spectro-temporal envelope.

Yet another popular abstraction is the *modulation spectrum* (Viemeister 1979; Dau et al 1997; Jepsen 2003; Joris et al 2004; Anden and Mallat 2014; Koumura et al 2019), defined as the spectrum of the *temporal envelope* within peripheral channels, possibly averaged over channels. It accounts for many aspects of sound, in particular speech (Elhilali et al 2003; Hermansky 2010) and, together with other statistics, captures the perceptually-relevant *texture* of many natural sounds (McDermott and Simoncelli 2011; McDermott et al 2013). Local fluctuations may serve also as cues to spectrally local features (Carney 2024). The quality of this abstraction is improved if the temporal envelope is first smoothed with a kernel tailored to the period of the sound (as described in the previous paragraph), to factor out periodicity-related fluctuations from slower perceptually-relevant modulations, thus avoiding interference between the two (Stein et al 2005).

Related to abstraction is *redundancy reduction*, and *summarization*. A benefit of detecting periodicity and estimating the period is that periodic segments can be cheaply represented by the waveform over one period, or its spectral equivalent (Fig. 5C). The ICC model contributes by detecting such periodic segments and their boundaries, and estimating the periods. This combination of *segmentation* and summarization may be seen as an initial layer in a hierarchical memory representation (de Cheveigné 2025).

This section hinted at various ways in which ICC *might* contribute to the elaboration of abstract representations of sound that *might* contribute to some of the top-down prediction schemes reviewed earlier. These ideas are presented schematically, obviously a more precise formulation is required to take them seriously as models of sensory processing.

## V. Discussion

This paper presented the low-level auditory processing model ICC as a form of *bottom-up* predictive coding. However, much of the appeal of the concept of predictive coding is specific to its *top-down* flavor; a reader who expected that flavor might feel short-changed. For this reason, I also investigated how ICC might relate to, or contribute to top-down predictive coding. At this point, the following hypothetical roles emerge:

1. The ICC model contributes an early link to the chain of abstractions postulated by most (all?) predictive coding models. One cannot achieve a hierarchy of abstraction if the output of each stage belongs to the same concrete space as its input, as is the case if each level sends a residual to the next. In contrast, the *parameters* of a model fit to a concrete pattern might be predicted, but this requires that that model (e.g., ICC) is defined.
2. The ICC model contributes to acoustic scene analysis, by its ability to suppress sounds with a harmonic, quasi-harmonic, or sparse spectrum. In this way, it might contribute to a “reverse acoustics” model on the lines of Kawato et al (1993).
3. The ICC model transforms input patterns that belong to the class of periodic sounds in such a way that they are distributed around zero, plausibly according to a Gaussian distribution. In this, it might play the role of a “Laplacifier” transform so that a predictive coding model on the lines of Friston (2009) meets its assumptions (Sect. IV, A).
4. The parameters of the ICC model (delays *τ*_*k*_) might be under top-down control. For example, a musical context might prime a listener as to the next pitch, or a decision to listen to source might translate into top-down imposition of delays that suppress competing sources. Multiple top-down parameters might be tried in sequence (or in parallel) to find one for which ICC best “fits” the input.
5. If predictive coding is reinterpreted as forming a compact statistical model of the stream of observations, ICC can contribute to the initial “chunking” by detecting intervals of quasi-periodicity and their boundaries, and by removing redundancy within such intervals. The magnitude of the ICC output serves as a measure of surprisal relative to the assumption of periodicity.

A stumbling block is the notion of prediction error as a *difference*, or *residual*. It would seem that Mumford’s (1992) warning about the need for translation between levels of abstraction was not heeded (including by Mumford himself). Objections to that notion are logical, as reviewed earlier, and also empirical. In a recent study, Furutachi et al (2024; Lowet and Uchida 2024) probed responses of putative “error neurons” in mouse primary visual cortex. With conditions carefully designed to test the difference hypothesis, they found that a surprising observation elicited an *enhanced* stimulus-specific response, rather than a difference. It is plausible that Mumford and others and others were aware of the issue, but that its implications got “lost in translation” when trying to make the model intelligible (as in Fig. 1C).

A prime motivation of top-down predictive coding is the need to explain efferent pathways within the brain. From the roles just enumerated it is not clear why they would need to be massive. One might speculate that massive efferent pathways are required by the need to propose, at level *n* + 1, a large number of alternative abstract explanations to be tested at level *n*.

Focusing on the auditory brainstem, the working hypothesis of a *single* delayed feedforward pathway within a channel might be over-restrictive. More plausible is a combination of multiple, within-channel predictors (as in LPC), possibly complemented by cross-channel interactions (e.g. Shamma 1984; Nelson and Carney 2004; de Cheveigné and Pressnitzer 2006; Carney 2002, 2024) and binaural interactions (Durlach 1964; Culling and Summerfield 1995; Beiderbeck et al 2018; Franken et al 2021). The multiple parameters involved (weights, delays) might help account for multiple efferent pathways.

An explanatory, reactive, or compressive model can better be fit and updated based on a buffer that extends into the past (Schmidhuber 2009). Predictive coding may contribute to *memory* of past observations (as opposed to a model of causes, decisions or actions) by helping to *segment* the input stream, and more generally helping to code observations in terms of already-acquired statistics, or internal explanatory models of the data. Rather than the message-passing hierarchy of Fig. 1C, memory would form a hierarchy of increasingly abstract statistics (de Cheveigné 2025). “Prediction error” might take one of the following forms: (1) a simple binary flag (fit vs no-fit) or a scalar (surprise), (2) a concise account of how the observation can be made to fit (e.g., “apply delay *τ* to the ICC model”), (3) a more exhaustive record or summary of the misfitting observations, “memorized” for future resolution.

To summarize, the simple bottom-up ICC model of auditory brainstem processing served as a pretext to review issues related to top-down predictive coding, highlighting the conceptual difficulties involved, and outlining approaches to dealing with them.

## Conclusion

The in-channel cancellation model describes a hypothetical form of low-level processing within the auditory brainstem, that offers a text-book example of predictive coding of the feedforward, bottom-up flavor, similar to earlier proposals for retinal processing (Srivinasan et al 1982). This paper briefly reviewed the model and its relations to bottom-up predictive coding, and examined how it might form part of an early layer within a predictive coding model of the feed-back, top-down flavor. Specifically, its segregation properties might contribute to a bottom-up “analysis” pathway that works hand-in-hand with a top-down predictive “synthesis” pathway (Kawato et al 1993). It could also participate to an early stage of statistical chunking of the sensory input that reorganizes it for more effective predictive coding. It might contribute to transform a non-Gaussian distribution of sensory regularities to better fit the Gaussian approximation assumed by many models of predictive coding.

## Acknowledgments

This work was supported by grants ANR-10-LABX-0087 IEC, ANR-10-IDEX-0001-02 PSL, and ANR-17-EURE-0017. Malcolm Slaney made helpful comments on an early draft.

